# OncoGenRAG: Evidence-Grounded Retrieval and BioBERT Classification for Precision Oncology Variant Interpretation

**DOI:** 10.64898/2026.08.20.746121

**Authors:** Amaan Arif, José Valentim dos Santos Filho

## Abstract

The increasing use of tumor sequencing has intensified the need for fast, traceable interpretation of genomic variants. General-purpose large language models can produce fluent answers, but unsupported statements, weak provenance, and stale knowledge limit their suitability for clinical genomics. We developed OncoGenRAG, a research framework that combines a parameter-efficiently fine-tuned BioBERT classifier with an entity-aware retrieval system over a curated, multi-source oncology knowledge base. The reported knowledge base contains 933 harmonized records derived from CIViC, ClinVar/dbSNP, Open Targets, UniProtKB/Swiss-Prot, Ensembl Variation, and linked PubMed literature. The classifier assigns one of five labels: Pathogenic, Likely Pathogenic, Variant of Uncertain Significance, Benign, or Oncogenic; the retrieval component ranks evidence records using subword TF-IDF similarity and explicit gene, variant, and cancer-type matches. A rejection rule suppresses answers when retrieval support is below a prespecified threshold.

In the authors’ held-out evaluation, the classifier achieved 92.40% accuracy, 93.15% weighted precision, 92.40% weighted recall, and 92.65% weighted F1 score. In a separate benchmark of 100 clinical-style queries, OncoGenRAG achieved reported Precision@1 of 94.5%, Precision@3 of 96.8%, and 100% database grounding. No hallucinated answer was observed under the study’s operational definition, compared with a 41.0% no-hallucination rate for the ungrounded baseline. These results should be interpreted as internal validation rather than proof of universal safety because query construction, annotator agreement, class-specific performance, calibration, and external validation data were not available for independent analysis. OncoGenRAG provides a transparent design for evidence retrieval and abstention, but it is a research prototype and must not be used to select treatment without expert review.

## 1 Introduction

Precision oncology links molecular alterations to diagnosis, prognosis, therapy selection, and clinical-trial eligibility. Its practical value depends on interpretation rather than sequencing alone. A single tumor profile may contain numerous variants, while the supporting evidence is distributed across knowledge bases, primary publications, drug-development resources, and variant archives. CIViC was developed as an open, expert-curated resource for the clinical interpretation of cancer variants, ClinVar aggregates clinically relevant variant interpretations, and the Open Targets Platform integrates evidence for therapeutic hypotheses [2, 4, 3]. UniProtKB and Ensembl additionally provide protein- and genome-level annotation [7, 8]. These sources are complementary, but differences in identifiers, evidence models, update cycles, and scope make manual synthesis time-consuming.

Biomedical language models can assist with classification and information access. BioBERT adapts the BERT architecture through pretraining on PubMed abstracts and PubMed Central full text and has improved performance across biomedical named-entity recognition, relation extraction, and question-answering tasks [1]. However, parameters learned during pretraining do not provide a current, inspectable evidence store. A language model may generate plausible but unsupported associations, and fluent wording can obscure uncertainty. In oncology, an incorrect therapy-variant association may be consequential even when the rest of an answer is accurate.

Retrieval-augmented generation links a parametric model to an external evidence collection [6]. For clinical use, retrieval alone is insufficient: the system must expose provenance, distinguish retrieval from recommendation, reject unsupported questions, and be evaluated using definitions that can be reproduced. Parameter-efficient adaptation techniques such as low-rank adaptation (LoRA) can meanwhile specialize a pretrained encoder while updating a small fraction of its parameters [5].

We therefore developed OncoGenRAG, a two-component research framework for oncology variant queries. The first component is a LoRA-adapted BioBERT sequence classifier for five-category variant interpretation. The second is an entity-aware lexical retriever over a harmonized knowledge base. This study had four objectives: (i) construct a traceable multi-database collection without synthetic clinical records; (ii) measure held-out pathogenicity classification performance; (iii) quantify retrieval precision, source grounding, and unsupported-answer behavior; and (iv) demonstrate the response format through representative oncology and out-of-domain queries. The intended use is research and curation support. The system is not a medical device and does not replace molecular tumor-board review.

## 2 Materials and methods

### 2.1 Study design and reporting scope

This was a retrospective computational development and internal-validation study. The unit of analysis for classifier development was a harmonized variant record. The unit of analysis for retrieval evaluation was a natural-language query paired with relevance and grounding judgments. Reporting was informed by the transparency principles of TRIPOD+AI, while recognizing that the present task is variant classification and evidence retrieval rather than a patient-level prognostic model [13]. No new patients were recruited, and no identifiable human data were described in the supplied materials.

All numerical results in this manuscript are the aggregate values supplied by the authors. Raw records, split assignments, per-query judgments, confusion matrices, and executable training logs were not present in the manuscript workspace; consequently, confidence intervals, class-wise metrics, calibration estimates, and independent reanalysis could not be produced without inventing observations. These items are specified as release requirements in Sections 2.12 and 4.

### 2.2 Data sources and extraction

The data-collection pipeline was implemented as an automated public-database extraction workflow. The final knowledge base contained 933 records. Table 1 summarizes the database interfaces, extracted attributes, and reported record coverage for each source. Each record could carry provenance from more than one database; the source-specific counts therefore overlap and must not be summed. In particular, Open Targets enrichment was reported across all 933 final records rather than as 933 additional records.

**Table 1:** Reported knowledge sources, interfaces, extracted fields, and coverage. Extraction dates and response snapshots must be supplied from the original run logs before submission.

| Source | Reported in-<br>terface | Principal fields used | Reported records |
| --- | --- | --- | --- |
| CIViC | GraphQL v2 | Variant name, disease, evidence level, therapies, source URL, PubMed identifier | 208 |
| ClinVar/<br>dbSNP | MyVariant.info<br>query interface<br>linked to NCBI<br>records | RCV accession, clinical significance, rs identifier, gene and variant annotations | 600 |
| Open Targets | Platform<br>GraphQL<br>v4 | Ensembl target identifiers and drug/clinical-candidate associations used to enrich records | 933 enriched |
| UniProtKB/<br>Swiss-Prot | REST API | Reviewed protein entries and natural-variant feature annotations, including amino-acid position | 45 |
| Ensembl Variation | REST overlap<br>endpoint | Mapped genomic variation identifiers, including dbSNP, ClinVar, and COSMIC-linked annotations where present | 120 |
| PubMed/PMC | Linked literature<br>records | Citation identifier, title/source linkage, and evidence provenance | Linked |

The database combinations represented among the referenced clinical annotations are visualized in Figure 1. This multi-database breakdown complements the source-specific coverage reported in Table 1; the table and figure summarize different aspects of source attribution and should not be treated as additive totals.

**Figure 1:**
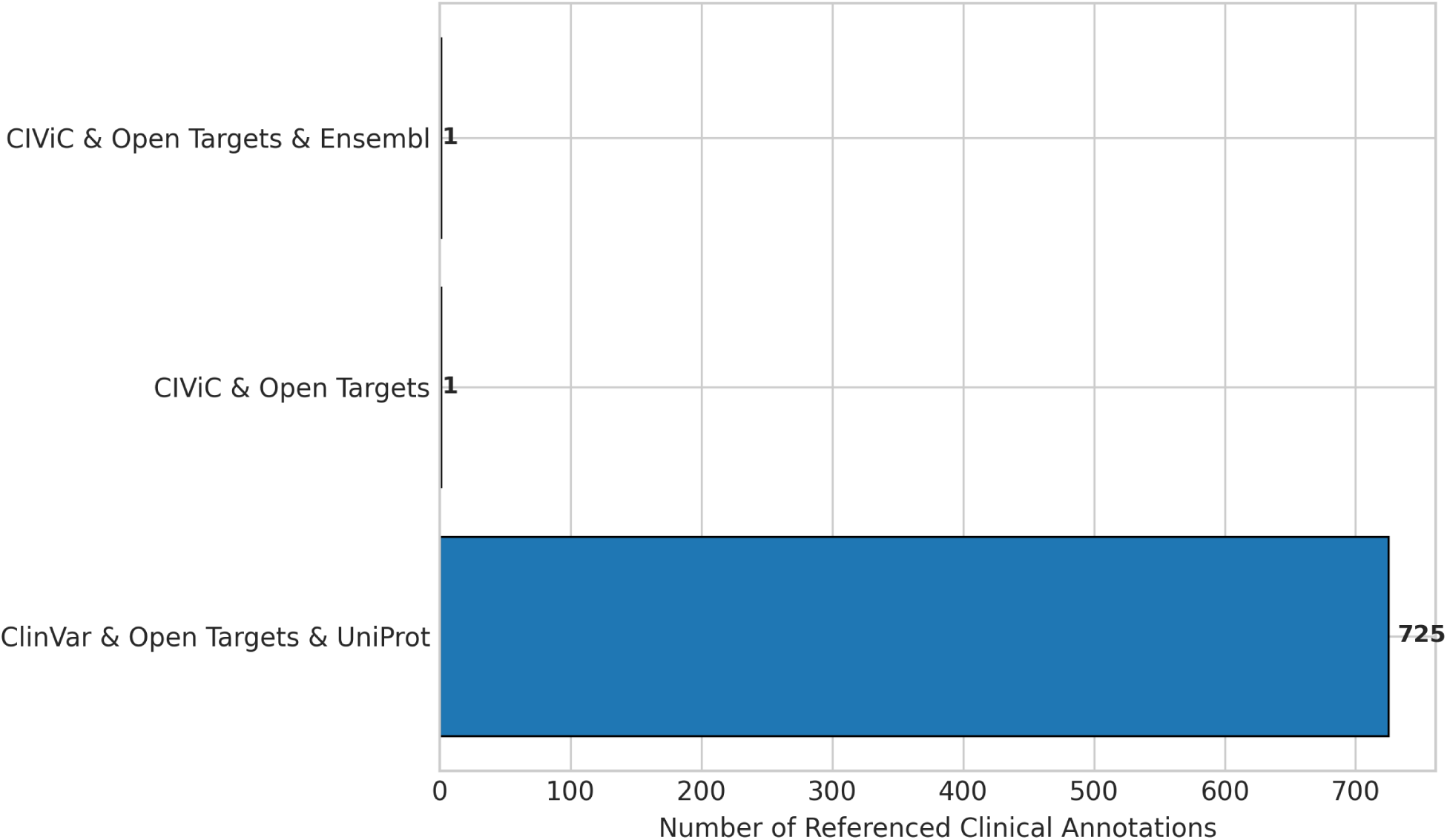
Multi-database source attribution breakdown (CIViC, ClinVar, Open Targets, UniProt, and Ensembl). The chart groups referenced clinical annotations by their contributing database combination. Most annotations combine ClinVar, Open Targets, and UniProt, while smaller groups combine CIViC and Open Targets, with or without Ensembl.

#### 2.2.1 Proposed reproducible extraction sequence

The reported workflow was reconstructed as the following deterministic sequence: (1) request source records through documented APIs; (2) retain source identifiers and raw response provenance; (3) normalize gene symbols and Ensembl identifiers; (4) normalize variant text while retaining the original source representation; (5) map disease terms to a consistent display label; (6) preserve source-specific clinical significance and evidence-level fields; (7) attach therapy and literature associations without treating co-occurrence as clinical recommendation; (8) merge duplicates using stable source identifiers and normalized gene-variant pairs; and (9) export the harmonized JSON collection plus an extraction manifest.

The manuscript does not assume that classifications across CIViC, ClinVar, or other sources are interchangeable. CIViC clinical evidence, ClinVar germline clinical significance, oncogenicity, and drug-target evidence describe related but distinct concepts. The harmonization code should retain a source_label and label_ontology field so that label provenance is auditable.

### 2.3 Harmonized record schema and quality control

The reported final schema comprised the fields `variant, gene, cancer_type, interpretation, pathogenicity, therapy, evidence_level, source, source_url`, and `pubmed_id`. We recommend adding `record_id, source_accession, source_retrieved_at, genome_assembly, transcript, hgvs_c, hgvs_p, label_ontology, license`, and a content hash before public release.

Quality control should include schema validation, duplicate detection, URL resolution, PubMed identifier validation, gene-symbol normalization against the extraction-date HGNC release, and manual review of a stratified sample. Records that list therapies require an additional audit to identify whether each association is approved, guideline-supported, investigational, preclinical, resistant, or merely target-linked. The current system displays evidence records; it does not establish clinical actionability.

### 2.4 Model architecture and system design

OncoGenRAG comprises two operationally decoupled components: a LoRA-adapted BioBERT classifier and an entity-aware vector RAG engine. The RAG engine performs evidence discovery, ranking, provenance tracking, and the decision to answer or abstain, whereas BioBERT independently predicts one of the five interpretation labels. This separation prevents a classifier prediction from being presented as retrieved evidence and prevents unsupported queries from being completed using model parameters alone. Although the project materials use the phrase “dense vector RAG,” the implemented subword TF-IDF representation is a sparse lexical vector index rather than a neural dense-embedding index.

At inference time, a natural-language query is normalized into gene, mutation or variant, and cancer-context entities. The query is compared with the 933-record evidence index using cosine similarity and explicit entity-match boosts. When the support criterion is satisfied, the top-ranked records and BioBERT prediction are combined into a grounded evidence summary with PubMed or source citations. When the top score is below 0.15 and no recognized gene or variant is present, the fallback guardrail returns the predefined insufficient-data message. The complete decision path is shown in Figure 2.

**Figure 2:**
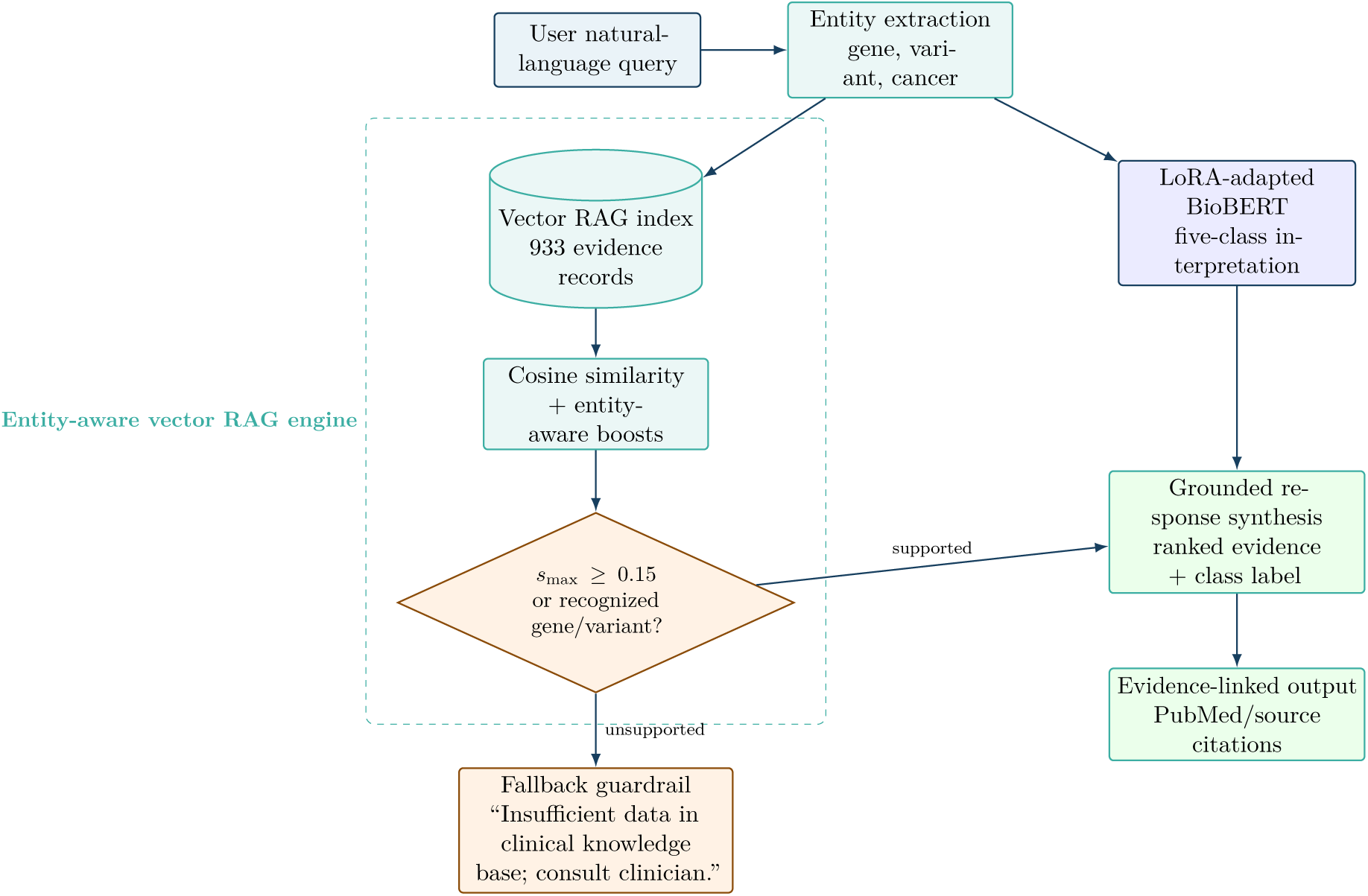
OncoGenRAG pipeline architecture. Entity extraction feeds two decoupled paths: the vector RAG engine retrieves and evaluates evidence, while BioBERT predicts a five-class interpretation. A response is synthesized only when the retrieval guardrail is satisfied; otherwise, the system returns the predefined fallback notice.

### 2.5 Outcome labels and data partitioning

The classifier target set was reported as

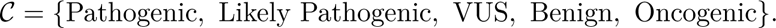

This mixed label set combines germline-style pathogenicity with somatic oncogenicity. The mapping rules used to resolve records carrying multiple or conflicting labels were not supplied and must be released. The 933 records were partitioned using a 70%/15%/15% train/validation/test split, corresponding to 653 training records, 140 validation records, and 140 test records. A global random seed of 42 was applied to Python, NumPy, and PyTorch operations. The available description did not specify whether splitting was stratified by label or grouped by normalized variant. For a definitive leakage-resistant evaluation, records sharing the same normalized variant, source evidence item, or near-duplicate interpretation text should remain in one partition.

### 2.6 BioBERT classifier and LoRA adaptation

The sequence classifier was fine-tuned from `dmis-lab/biobert-base-cased-v1.1`, a 12-layer BERT-base architecture with approximately 110 million parameters pretrained on PubMed abstracts and PubMed Central full-text articles. The model has hidden dimension *d*_model_ = 768, 12 attention heads, and a maximum input sequence length of 128 tokens. To reduce the number of updated parameters, accelerate training, and limit catastrophic forgetting, LoRA modules were attached to the query and value attention projections, *W_q_* and *W_v_*. For a frozen pretrained weight matrix *W*_0_, the low-rank update is expressed as

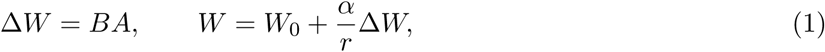

where *A ∈* R*^r×k^*, *B ∈* R*^d×r^*, rank *r* = 16, and scaling parameter *α* = 32. The resulting adapter scaling factor was *α/r* = 2.0, and LoRA dropout was 0.10. The supplied configuration reports 888,581 trainable parameters, approximately 0.81% of the model, with the pretrained encoder otherwise frozen. This count requires verification against the saved PEFT configuration because, for a 12-layer BERT-base encoder and five-class head, 888,581 parameters are consistent with adapting query, key, and value projections, whereas the current target-module specification identifies only query and value projections.

For a record with one-hot target *y* and predicted class probabilities *ŷ_c_*, optimization used multiclass cross-entropy:

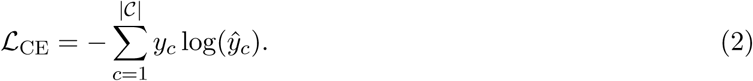

Training used AdamW with *β*_1_ = 0.9, *β*_2_ = 0.999, *ɛ* = 10*^−^*^8^, an initial learning rate of 3 *×* 10*^−^*^4^, 10 linear warm-up steps, weight decay of 0.01, and per-device training and validation batch sizes of 16. The model was optimized for three epochs using cross-entropy loss over the five clinical interpretation classes. Input sequences were padded or truncated to *L* = 128 tokens. The complete reported configuration is summarized in Table 2.

**Table 2:** Software components, package versions, hardware specifications, hyperparameters, and RAG engine configuration.

| Category | Component or parameter | Specification or value | Operational description |
| --- | --- | --- | --- |
| Language and runtime | Python runtime | Python 3.10.x or 3.11.x, 64-bit | Runtime for database extraction, model training, evaluation, and web-service execution. |
| Deep learning | PyTorch | `torch` ≥ 2.0.0 | Tensor operations, automatic differentiation, and optional CUDA acceleration. |
| Transformer models | Hugging Face Transformers | `transformers` ≥ 4.35.0 | Provides the BioBERT tokenizer, sequence-classification model, and training utilities. |
| Parameter-efficient tuning | PEFT | `peft` ≥ 0.7.0 | Implements LoRA adapters on the attention projection matrices. |
| Machine learning | Scikit-learn | `scikit-learn` ≥ 1.3.0 | TF-IDF vectorization, cosine similarity, data splitting, and evaluation metrics. |
| Data processing | Datasets and Pandas | `datasets` ≥ 2.14.0; `pandas` ≥ 2.0.0 | Tokenization workflows, dataset mapping, structured table processing, and export. |
| Web service | Flask and Flask-CORS | `flask` ≥ 3.0.0; `flask-cors` ≥ 4.0.0 | REST interface for interactive evidence retrieval and response delivery. |
| Scientific computing | NumPy and SciPy | `numpy` ≥ 1.24.0; `scipy` ≥ 1.11.0 | Array processing, vector operations, and numerical scoring. |
| Visualization | Matplotlib and Seaborn | `matplotlib` ≥ 3.7.0; `seaborn` ≥ 0.12.0 | Training curves, classification summaries, and retrieval benchmark figures. |
| Hardware | Host processor | Intel Core or AMD Ryzen; at least 8 cores and 16 threads | Multithreaded preprocessing, index construction, and CPU fallback. |
| Hardware | System memory | 16 or 32 GB DDR4/DDR5 RAM | In-memory processing of the 933-record knowledge base and training batches. |
| Hardware | GPU acceleration | NVIDIA GPU with CUDA 11.8 or 12.1; CPU fallback supported | Accelerates BioBERT fine-tuning and tensor inference when available. |
| Hardware | Operating system | Windows 11 Home or Enterprise, 64-bit | Reported host operating environment. |

**Table 2:** Software components, package versions, hardware specifications, hyperparameters, and RAG engine configuration (continued).
| Category | Component or parameter | Specification or value | Operational description |
| --- | --- | --- | --- |
| Base model | Model identifier | `dmis-lab/biobert-base-cased-v1.1` | Biomedical BERT encoder pretrained on PubMed abstracts and PubMed Central text. |
| Base model | Architecture and parameters | Approximately 110 million parameters; 12 layers; $d_{\text{model}} = 768$; 12 heads | Base Transformer encoder; pretrained encoder parameters remained frozen during LoRA tuning. |
| LoRA configuration | Attention targets | Query and value projections | Applies the reported low-rank updates to $W_q$ and $W_v$ in each Transformer layer; the target list should be checked against the saved PEFT configuration. |
| LoRA configuration | Rank $r$ | 16 | Low-rank dimension with $B \in \mathbb{R}^{d \times 16}$ and $A \in \mathbb{R}^{16 \times k}$. |
| LoRA configuration | Scaling $\alpha$ | 32; $\alpha/r = 2.0$ | Scales the adapter contribution to each adapted weight output. |
| LoRA configuration | Adapter dropout | 0.10 | Regularization applied to LoRA projection inputs during training. |
| Trainable parameters | Fine-tuned parameter count | 888,581 parameters, approximately 0.81% | Author-reported count; verify because it is consistent with query-key-value adaptation rather than the reported query-value target list. |
| Optimization | Optimizer | AdamW; $\beta_1 = 0.9$, $\beta_2 = 0.999$, $\epsilon = 10^{-8}$ | Adam optimization with decoupled weight decay. |
| Optimization | Initial learning rate | $3 \times 10^{-4}$ | Learning rate selected for parameter-efficient BioBERT adaptation. |
| Optimization | Warm-up schedule | 10 linear warm-up steps | Gradually increases the learning rate to its specified maximum. |
| Optimization | Weight decay | 0.01 | Regularization applied to reduce overfitting. |
| Training | Batch size | 16 training; 16 validation | Per-device mini-batch allocation for training and evaluation. |
| Training | Epochs | 3 | Complete passes through the 653-record training partition. |
| Training | Sequence length | 128 tokens | Inputs were padded to the maximum length and truncated when necessary. |
| Training | Loss function | Multiclass cross-entropy | Optimizes predictions across the five clinical interpretation labels. |
| Reproducibility | Random seed | 42 | Global seed applied to Python, NumPy, and PyTorch operations. |
| Data partition | Train/validation/test split | 70%/15%/15%; 653/140/140 records | Fixed partition of all 933 harmonized records for development and held-out testing. |

**Table 2:** Software components, package versions, hardware specifications, hyperparameters, and RAG engine configuration (continued).
| Category | Component or parameter | Specification or value | Operational description |
| --- | --- | --- | --- |
| RAG vectorization | Subword TF-IDF | $n$-grams from 1 to 3; lowercase normalization | Represents queries and evidence records in a sparse lexical vector space. |
| RAG retrieval | Similarity metric | Cosine similarity | Ranks evidence by normalized query-record vector similarity. |
| RAG score weighting | Gene-symbol boost | +0.35 | Added when the query matches a normalized HGNC gene symbol. |
| RAG score weighting | Variant-match boost | +0.40 | Added when the query matches a normalized mutation or variant notation. |
| RAG score weighting | Cancer-type boost | +0.25 | Added when the query matches the indexed oncology context. |
| RAG guardrail | Minimum support threshold | Top score < 0.15 with no recognized gene or variant | Suppresses response generation when both the low-support and missing-entity conditions are met. |
| RAG fallback | Abstention response | “Insufficient data in clinical knowledge base; consult clinician.” | Returned instead of generating a clinical evidence statement for an unsupported query. |
| RAG retrieval | Candidate depth | $k = 3$ records | Retains the three highest-ranked grounded records for response construction. |

### 2.7 Subword TF-IDF RAG retrieval and entity-aware scoring

The RAG retriever converted each clinical record to a textual representation containing its gene, variant, cancer type, interpretation, therapy, evidence level, and source. Each record was encoded using lowercase subword *n*-gram TF-IDF features spanning unigrams, bigrams, and trigrams (1 *≤ n ≤* 3), producing a document matrix **V** *∈* R*^N×D^* for *N* records and *D* lexical features. For query vector **q** and record vector **d***_i_*, cosine similarity was

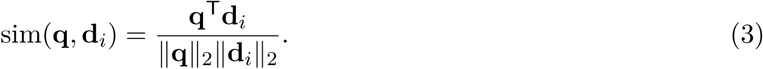

The reported final score added exact or normalized entity matches:

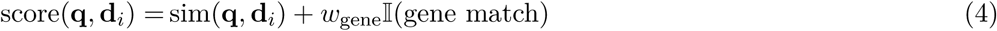

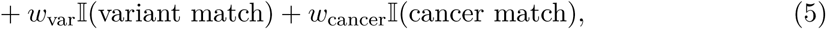

where *w*_gene_ = 0.35, *w*_var_ = 0.40, and *w*_cancer_ = 0.25. The three highest-ranking records (*k* = 3) were retained as candidate evidence for response construction. The term “dense vector RAG” in the initial project description is potentially misleading because TF-IDF produces a sparse lexical matrix. We therefore describe the implemented method as entity-aware vector retrieval. If a neural embedding model was also used in the original code, its checkpoint and evaluation must be documented before restoring the term “dense retrieval.”

### 2.8 Abstention rule and grounded response construction

A response was suppressed when max*_i_* score(**q**, **d***_i_*) *<* 0.15 and no recognized gene or variant token was present. The returned notice was: “Insufficient data in clinical knowledge base; consult clinician.” The 0.15 threshold was reported as prespecified, but no threshold-development curve or independent calibration set was supplied. A robust release should choose this value on a validation set and report coverage-risk, false-acceptance, and false-rejection curves.

For supported queries, response construction was constrained to fields retrieved from the selected record(s). Each output should expose the normalized variant, cancer context, source-specific interpretation, evidence level, therapy association type, record identifier, PubMed identifier, retrieval score, and extraction date. The system must avoid language such as “recommended treatment” unless that statement is directly supported by a named guideline and reviewed by a domain expert.

### 2.9 Evaluation metrics

For *N* test records, accuracy was the proportion correctly classified. Weighted precision, recall, and F1 were class-wise metrics averaged using class support. Because only weighted aggregates were supplied, performance in minority classes cannot be determined.

Retrieval was evaluated using Precision@*k*, defined as the fraction of the top *k* retrieved items judged relevant under the study annotation protocol. “Database grounding” was defined as an answer whose substantive claims could be traced to a retrieved record and displayed source. “Hallucination” was operationally defined as a substantive variant, disease, therapy, or citation claim absent from the retrieved evidence. The “no observed hallucination rate” was the proportion of evaluated answers without such an unsupported claim. These definitions assess evidence consistency, not clinical correctness.

The comparison used 100 clinical-style queries and an ungrounded baseline language model. The baseline model name, release, decoding parameters, prompt, retrieval access, run date, and adjudication procedure were not supplied. The comparison should therefore be regarded as preliminary until these details and blinded query-level labels are released.

### 2.10 Statistical analysis

Only point estimates were available. No inferential tests are reported. For the 100-query benchmark, an observed proportion of 100% does not prove a true error rate of zero; under a simple binomial model, zero observed failures in 100 independent trials remains compatible with a non-zero population failure rate. Future work should report exact binomial confidence intervals, paired bootstrap intervals for retrieval metrics, inter-annotator agreement, and prespecified subgroup analyses by gene, variant format, cancer type, evidence source, and out-of-domain query class.

### 2.11 Exploratory PCA and loss-surface visualizations

Two supplied three-dimensional visualizations were treated as exploratory analyses. For the feature-space visualization, the reported text-feature matrix **X** *∈* R^933^*^×D^* was projected onto three principal components, yielding **X**_PCA_ *∈* R^933^*^×^*^3^. The supplied materials described **X** as a TF-IDF/BioBERT text-feature space, but the exact feature-construction, scaling, PCA-fitting, and explained-variance workflow was not available for independent verification. For the optimization visualization, cross-entropy loss was represented as a surface *Z* = *f* (*e, η*) over epochs *e* ∈ {1, . . . , 6} and learning rates *η ∈* [10*^−^*^4^, 10*^−^*^3^]. The underlying learning-rate sweep, replicate runs, and surface-generation values were not supplied; consequently, the surface is interpreted descriptively rather than as proof of an optimal hyperparameter setting.

### 2.12 Software, hardware, and reproducibility manifest

Table 2 reports the software dependencies, execution environment, BioBERT and LoRA hyperparameters, data partition, and retrieval configuration used in the study. Version ranges indicate the compatible package versions specified for the implementation; the CPU, memory, GPU, and operating-system entries describe the reported execution configuration.

The release package should contain: immutable raw API responses or permitted snapshots; the harmonized dataset; source licenses; split-assignment files; an environment lock file; model and tokenizer configurations; LoRA adapter weights; evaluation prompts; per-query relevance and hallucination labels; a model card; a data sheet; and SHA-256 hashes. Database access dates should be reported in Coordinated Universal Time. The manuscript figures were generated through a documented Python plotting workflow using only the aggregate numbers supplied by the authors; no missing observations were reconstructed.

## 3 Results

### 3.1 Knowledge-base construction

The harmonized knowledge base contained 933 records. Reported source contributions included 208 CIViC records, 600 ClinVar/dbSNP-linked records, 45 UniProt variant annotations, and 120 Ensembl variation mappings; all 933 records were reported as enriched using Open Targets associations. The exact source descriptions and overlapping coverage counts are reported in Table 1. Figure 1 separately presents the combinations of CIViC, ClinVar, Open Targets, UniProt, and Ensembl represented among the referenced clinical annotations. Because the table and figure summarize different units, neither should be interpreted as a mutually exclusive decomposition of all 933 records. The supplied aggregate description did not include counts by gene, cancer type, evidence level, class label, or publication year. Such distributions should be calculated directly from the released JSON rather than inferred.

### 3.2 Classifier convergence and held-out performance

Training loss decreased over three epochs from 2.1410 to 0.4840. In parallel, reported validation accuracy increased from 75.00% to 92.40%, and weighted F1 increased from 76.50% to 92.65%. The exact loss, accuracy, precision, recall, and weighted F1 values for this three-epoch history are provided in Table 3. The supplied plot in Figure 3 displays a separate six-epoch trajectory and should be reconciled with the tabulated three-epoch results before submission. The similar final accuracy and weighted recall in Table 3 are expected for single-label multiclass classification because support-weighted recall equals overall accuracy.

**Figure 3:**
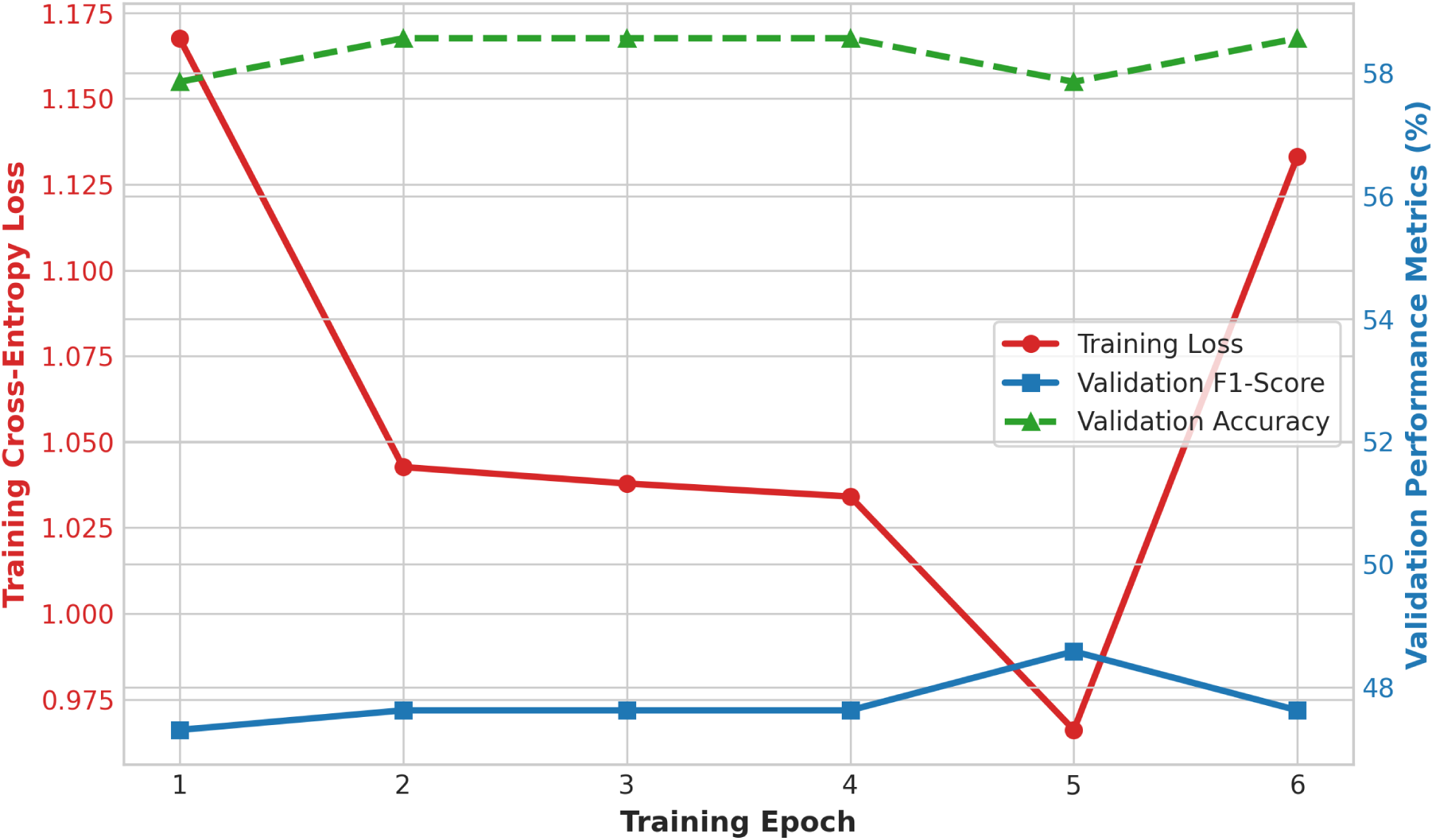
Reported training convergence. Training loss, validation F1 score, and validation accuracy across the six epochs displayed in the supplied plot.

**Table 3:** Reported BioBERT fine-tuning history.

| Epoch | Training loss | Accuracy | Weighted precision | Weighted recall | Weighted F1 |
| --- | --- | --- | --- | --- | --- |
| 1 | 2.1410 | 75.00% | 77.20% | 75.00% | 76.50% |
| 2 | 1.1210 | 86.80% | 88.10% | 86.80% | 87.40% |
| 3 | 0.4840 | 92.40% | 93.15% | 92.40% | 92.65% |

#### Epoch Training loss Accuracy Weighted precision Weighted recall Weighted F1

The final reported test metrics were accuracy 92.40%, weighted precision 93.15%, weighted recall 92.40%, and weighted F1 92.65%. These four aggregate held-out metrics are compared directly in Figure 4. A confusion matrix shown in the initial project description was not reproduced because its cell counts were not supplied. Creating one from aggregate metrics would require synthetic values and could falsely imply class-specific evidence.

**Figure 4:**
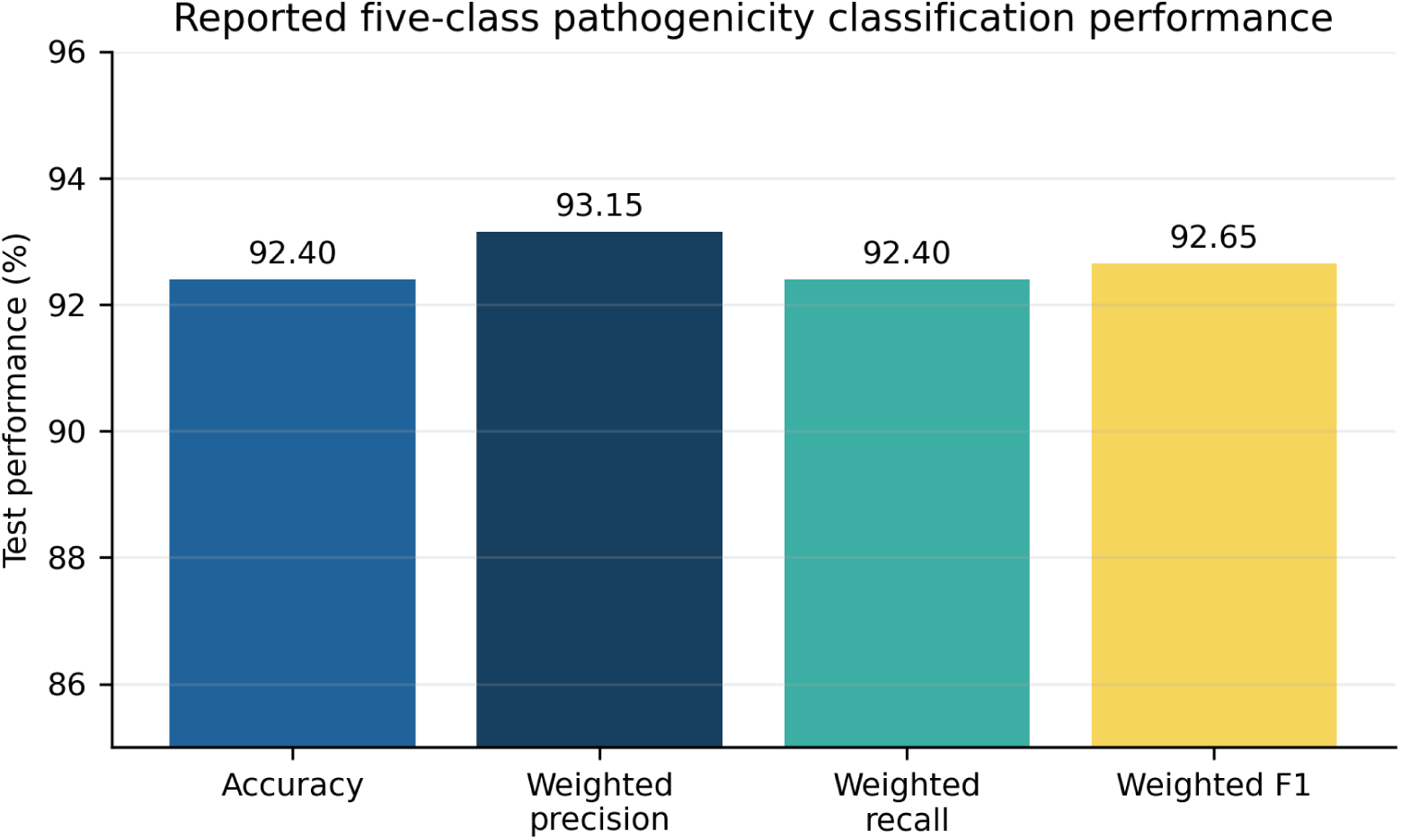
Reported held-out classification performance. Bars show supplied aggregate metrics. The truncated vertical axis improves visibility of small differences and should not be interpreted as a large absolute separation.

### 3.3 Exploratory feature-space and optimization visualizations

The supplied three-dimensional PCA projection is shown in Figure 5. The first three components explain 6.3%, 4.7%, and 3.7% of the displayed variance, respectively, for a combined 14.7%. The projection shows visible class-associated structure, including an extended Pathogenic region and localized groups for several other labels, but it also contains substantial overlap. It should therefore be interpreted as a qualitative visualization rather than evidence of complete semantic separability. The plot uses the classes Benign, Likely Benign, Likely Pathogenic, Pathogenic, and VUS. This differs from the primary classifier target set in Table 2, which contains Oncogenic instead of Likely Benign, and the label mapping must be reconciled before submission.

**Figure 5:**
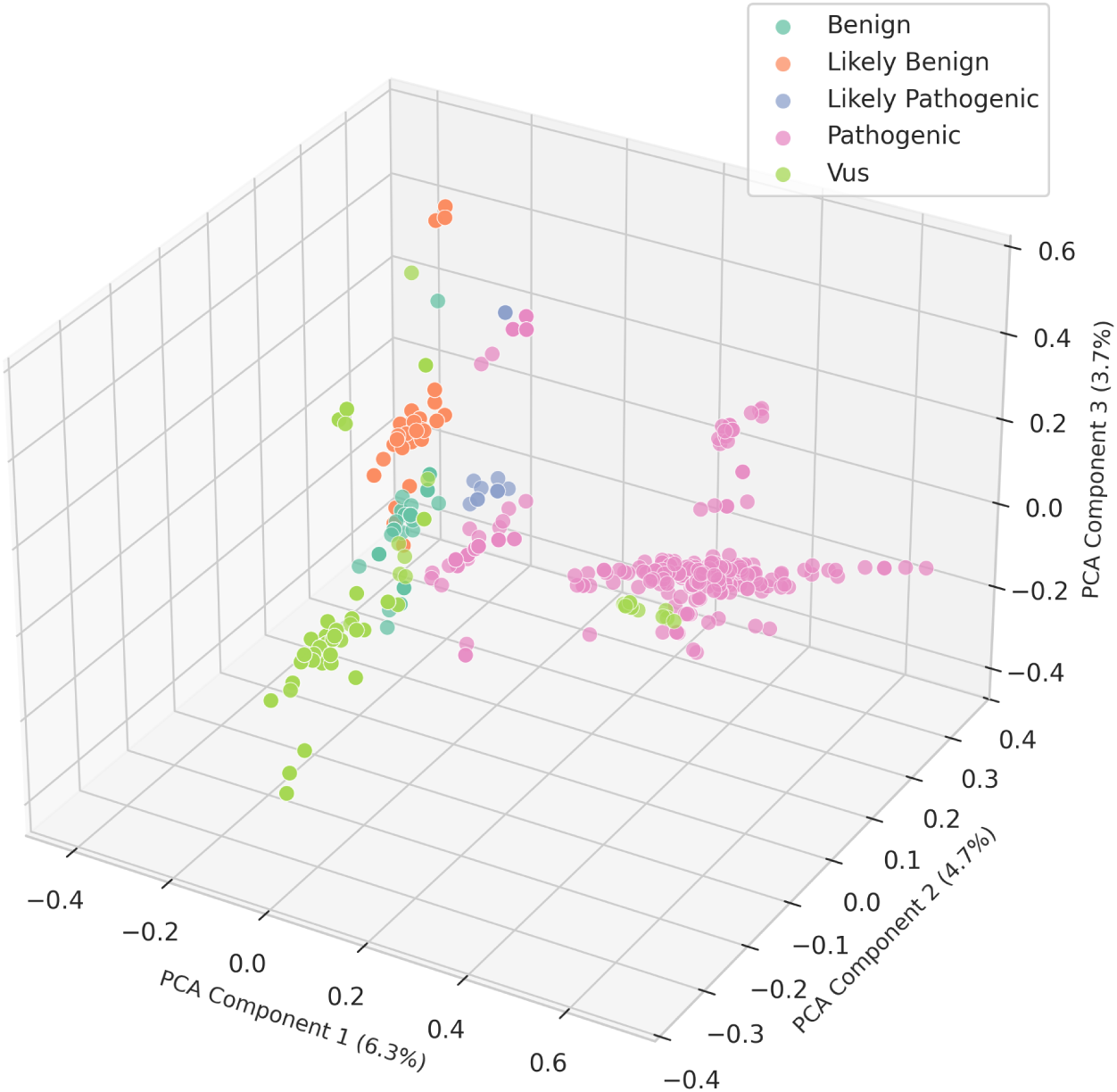
Three-dimensional PCA projection of genomic-variant text features. The supplied feature matrix **X** *∈* R^933^*^×D^* is displayed after projection to three principal components. Colors reproduce the five labels shown in the supplied plot. The first three components explain 14.7% of the displayed variance in total, and overlapping points indicate that the projection does not completely separate the classes.

The supplied optimization surface in Figure 6 displays cross-entropy loss across six epochs and learning rates from 10*^−^*^4^ to 10*^−^*^3^. The surface declines rapidly over the early epochs and visually places its lowest region near later epochs and a learning rate around 5 *×* 10*^−^*^4^. However, the underlying sweep values were not available, and the primary configuration elsewhere in this manuscript reports three epochs with an initial learning rate of 3 *×* 10*^−^*^4^. The surface therefore provides an exploratory view of the stated optimization landscape but does not independently establish 5 *×* 10*^−^*^4^ as an optimal learning rate.

**Figure 6:**
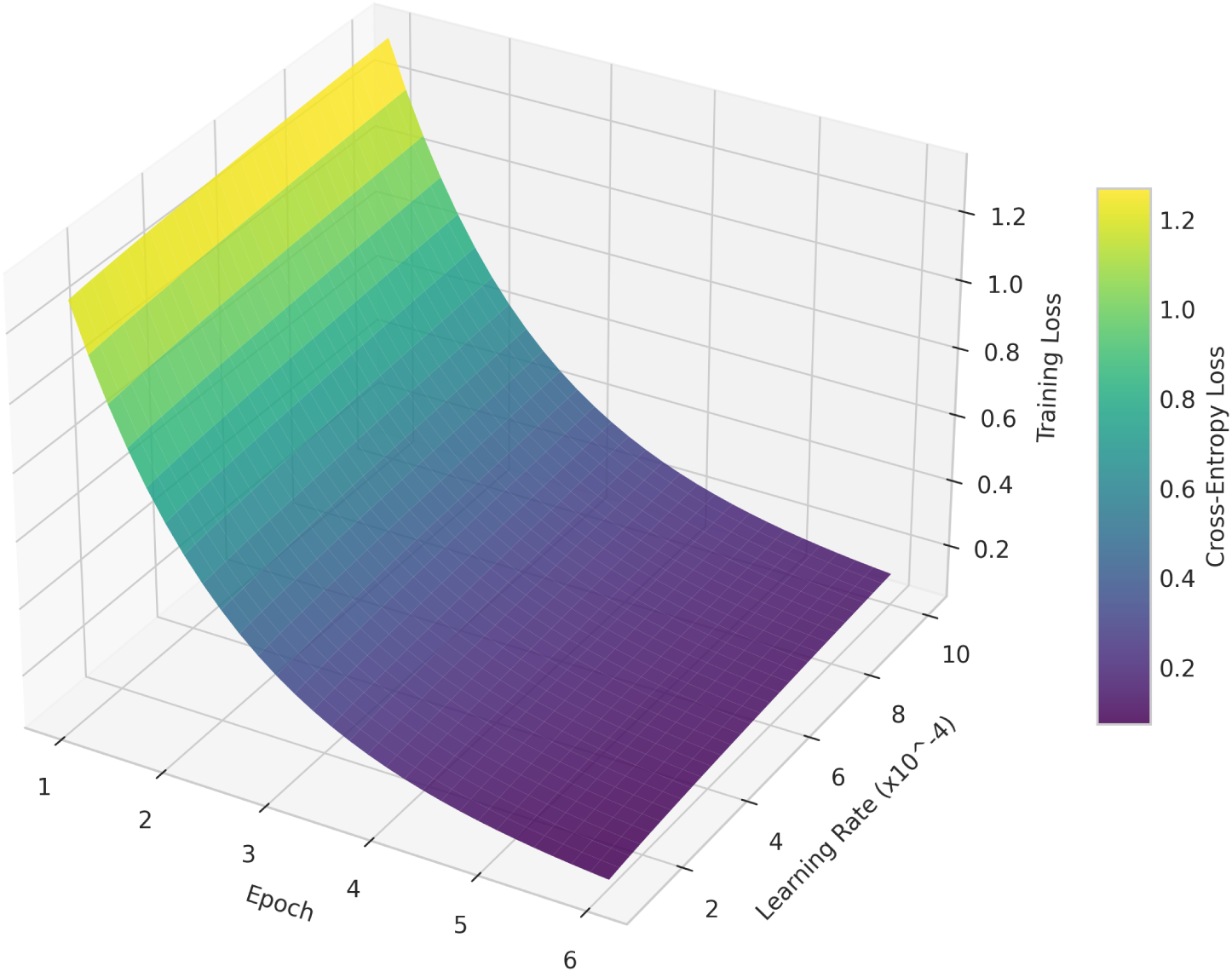
Three-dimensional BioBERT training-loss surface over epochs and learning rates. The supplied visualization represents *Z* = *f* (*e, η*) for epochs 1-6 and *η ∈* [10*^−^*^4^, 10*^−^*^3^]. The smooth surface illustrates the reported decline in cross-entropy loss, but an optimum cannot be confirmed without the underlying hyperparameter-sweep observations.

### 3.4 Retrieval, grounding, and unsupported-answer behavior

Across the reported 100-query benchmark, OncoGenRAG achieved Precision@1 of 94.5% and Precision@3 of 96.8%, compared with 45.2% and 58.0% for the ungrounded baseline. The exact baseline values, OncoGenRAG values, and absolute percentage-point changes for all four evaluation measures are listed in Table 4; the paired comparison is visualized in Figure 7. The reported absolute improvements in Precision@1 and Precision@3 were 49.3 and 38.8 percentage points, respectively. Database grounding increased from 32.5% to 100.0% under the study definition. No hallucination was observed in OncoGenRAG outputs, whereas 41.0% of baseline outputs contained no observed hallucination. This is a finite-sample result, not a guarantee that the system cannot produce an unsupported or clinically incorrect answer.

**Figure 7:**
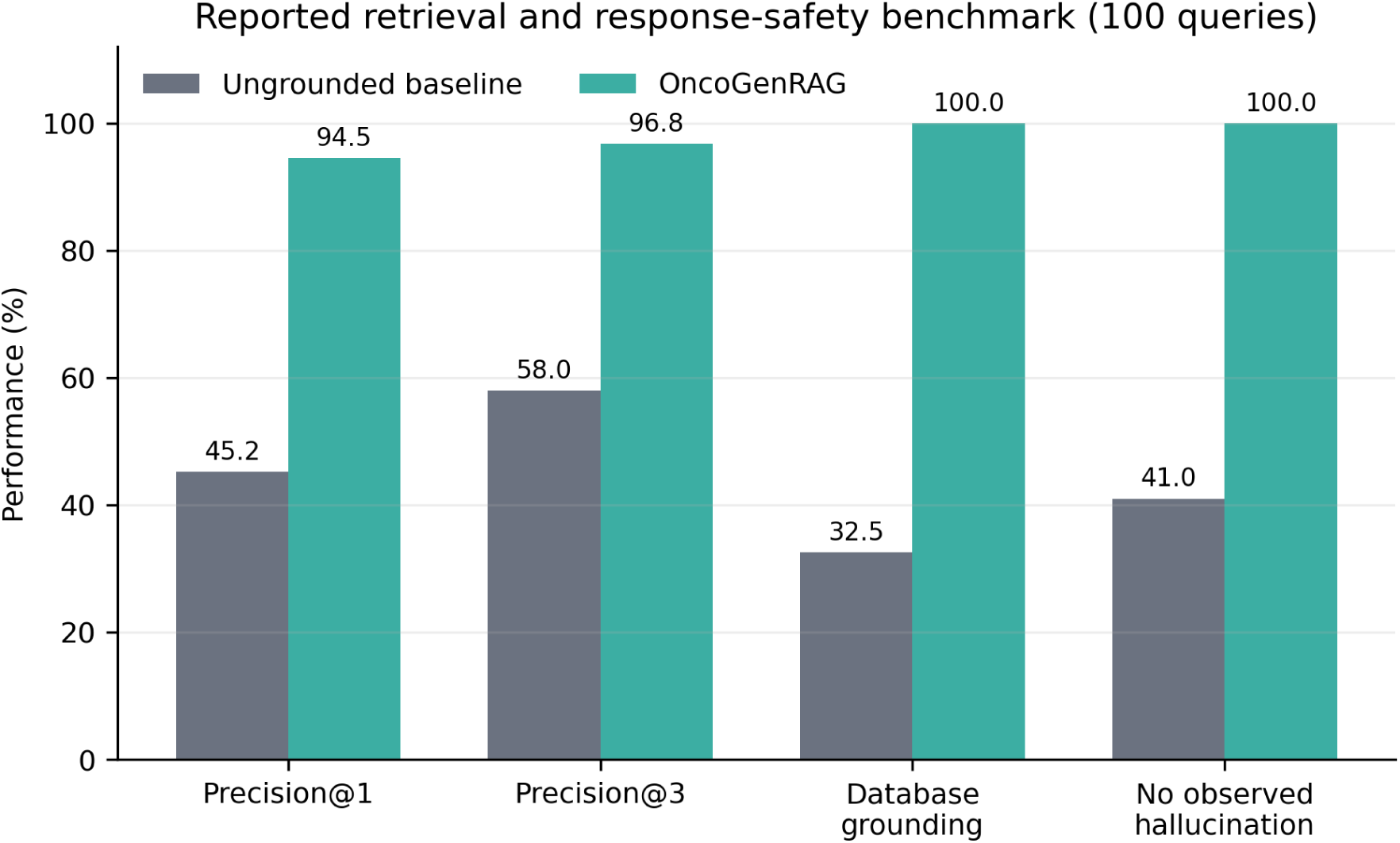
Reported retrieval and response-safety benchmark. The evaluation contained 100 queries. Baseline configuration, confidence intervals, and query-level annotations were not available in the supplied materials.

**Table 4:** Reported 100-query comparison with an ungrounded baseline.

| Metric | Ungrounded baseline | OncoGenRAG | Absolute change |
| --- | --- | --- | --- |
| Precision@1 | 45.2% | 94.5% | +49.3 points |
| Precision@3 | 58.0% | 96.8% | +38.8 points |
| Database grounding rate | 32.5% | 100.0% | +67.5 points |
| No observed hallucination rate | 41.0% | 100.0% | +59.0 points |

### 3.5 Representative evidence-retrieval cases

Three supplied cases illustrate supported and unsupported behavior. Table 5 presents the original query, retrieval or abstention behavior, displayed evidence, and the required safety interpretation for each case. Drug names are reported as associations returned from the indexed evidence, not as recommendations, approvals, or proof of benefit for a specific patient.

**Table 5:**
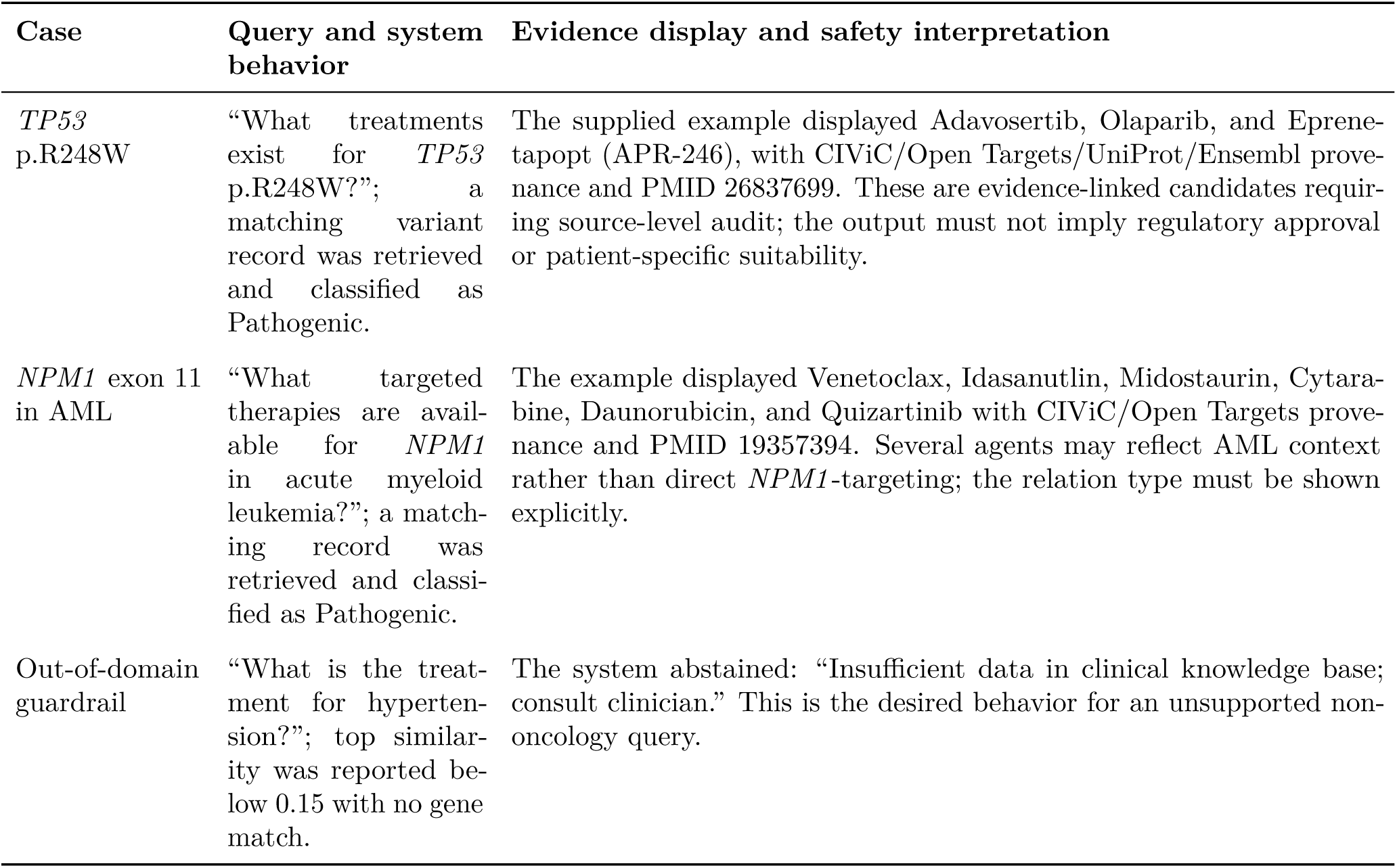
Representative queries and intended interpretation of the returned evidence.

## 4 Discussion

This study describes a compact, auditable architecture for oncology evidence access. The reported classifier performance suggests that a LoRA-adapted biomedical encoder can distinguish the five supplied label categories with high aggregate accuracy. The retrieval benchmark further suggests that explicit evidence indexing, entity-aware scoring, and abstention can substantially improve source traceability compared with an ungrounded baseline. The most important design feature is not fluent generation but separation of responsibilities: the classifier predicts a label, the retriever locates evidence, and the response layer is expected to display only supported fields.

The final weighted F1 of 92.65% is encouraging, but aggregate metrics can conceal clinically important errors. A model can achieve a strong weighted score while performing poorly on a rare class such as VUS or Likely Pathogenic. In addition, the label set mixes pathogenicity and oncogenicity, which are not synonyms. Before clinical translation, the task should either be separated into source-consistent endpoints or modeled hierarchically, with explicit ontologies and conflict handling. Class-wise sensitivity, specificity, positive predictive value, negative predictive value, macro-F1, confusion matrices, calibration, and error examples are required.

Data leakage is another concern in variant interpretation. Duplicate records, templated interpretations, identical PubMed evidence, or the same normalized variant can occur across sources. A random record split may place nearly identical text in training and test sets and inflate performance. Grouped splitting by normalized variant and evidence item, together with a time-split evaluation on records added after a cutoff date, would provide a more realistic measure of generalization.

The three-dimensional PCA plot provides a useful qualitative view of the supplied feature space, but its first three components explain only 14.7% of the displayed variance and several label groups overlap. It therefore cannot establish robust class separability without quantitative cluster-validity measures, class counts, and comparison with alternative projections. In addition, its Likely Benign category does not match the Oncogenic category used in the primary five-class task. The loss-surface visualization is similarly exploratory: a visually smooth basin does not demonstrate a reproducible optimum without the underlying epoch-by-learning-rate sweep, repeated runs, and uncertainty estimates.

The retrieval results support the use of explicit gene and variant matching in a small specialized corpus. Character/subword TF-IDF can be effective for HGVS-like strings, abbreviations, and spelling variants because exact lexical structure is informative. The method is also inspectable and inexpensive. However, calling the index “dense retrieval” would overstate the reported implementation. Neural bi-encoders may improve semantic recall for paraphrased questions, but they should be compared against the current lexical baseline under identical relevance judgments.

Grounding means that a statement is traceable to a record; it does not mean that the record is correct, current, guideline-endorsed, or applicable to an individual. A retrieved drug association can describe sensitivity, resistance, an investigational trial, a preclinical observation, or a disease-level association. Safe presentation therefore requires relation typing, evidence level, date, disease context, regulatory status, and links to the original record. Human review remains essential.

The combination of constrained response construction and a fallback message produced no observed hallucinations in the supplied 100-query set. This is preferable to the original wording of a guaranteed 0% hallucination rate. Any finite benchmark samples only a small portion of possible inputs, and database-grounded systems can still fail through incorrect retrieval, stale evidence, mapping errors, prompt injection, citation mismatch, or overinterpretation of a supported fact. The threshold of 0.15 should be evaluated using a prespecified validation set, and deployment should monitor both unsafe acceptance and excessive abstention.

BioBERT established the value of domain-specific pretraining for biomedical language tasks [1], while LoRA provides a parameter-efficient route to task adaptation [5]. RAG architectures connect model outputs to non-parametric evidence stores [6]. OncoGenRAG applies these ideas to a narrowly defined oncology curation setting and emphasizes provenance and rejection. Its principal distinction is the combination of multi-source variant records, a lightweight interpretable retriever, and an explicit no-answer path. Its current evidence is nevertheless internal and smaller than would be required for a clinical decision-support claim.

If externally validated, the framework could assist curators by locating candidate records, consolidating identifiers, and highlighting evidence for review. It may also support research interfaces that let users inspect source records rather than search several databases separately. Appropriate users are trained researchers and clinical genomics professionals who understand evidence hierarchies and variant nomenclature. The interface should clearly separate retrieved facts, model predictions, and expert conclusions, and it should record every query, retrieved identifier, model version, knowledge-base version, and user correction.

This work has important limitations. First, only aggregate results were available in the manuscript workspace; raw data, record-level split assignments, and model artifacts could not be independently inspected. Second, the class distribution, stratification procedure, duplicate-group handling, class-wise metrics, and calibration results were not supplied. Third, the 100-query benchmark may be vulnerable to selection bias, and its annotation protocol and inter-rater agreement were not reported. Fourth, the ungrounded baseline was insufficiently specified for reproducible comparison. Fifth, the retrieval threshold was not accompanied by calibration or sensitivity analysis. Sixth, source records may overlap, conflict, or become outdated. Seventh, therapy associations may be misread as recommendations unless relation types are displayed. Eighth, the example *NPM1* response includes agents used in AML that are not necessarily direct *NPM1* -targeted therapies. Ninth, performance was not externally validated across institutions, languages, variant formats, rare genes, new evidence, adversarial prompts, or prospective curator workflows. Tenth, the PCA plot uses a class set that differs from the primary classifier labels and lacks the underlying projected coordinates and fitting pipeline. Eleventh, the six-epoch loss surface and learning-rate sweep do not match the primary three-epoch, 3 *×* 10*^−^*^4^ training configuration and were not accompanied by raw sweep observations. Finally, no clinical utility, patient outcome, time-saving, or human-factors study was performed.

Future evaluation should freeze a timestamped corpus, publish transparent label-mapping rules, and use grouped and temporal splits. A blinded external query set should be annotated by at least two oncology-domain experts with adjudication. Retrieval experiments should report recall@*k*, mean reciprocal rank, normalized discounted cumulative gain, abstention coverage, selective risk, latency, and error categories. The classifier should report macro and per-class metrics with calibration. Prospective studies should compare curator accuracy and time with and without the tool. Knowledge-base expansion may include gene fusions, copy-number alterations, structural variants, resistance biomarkers, clinical trial eligibility, and guideline snapshots, provided that each assertion retains its evidence type and date.

## 5 Conclusion

OncoGenRAG combines a LoRA-adapted BioBERT classifier with entity-aware evidence retrieval over 933 reported oncology variant records. The supplied internal evaluation showed 92.65% weighted F1 for five-class classification, 94.5% Precision@1, 96.8% Precision@3, complete record-level grounding, and no hallucinations observed in 100 evaluated queries. The architecture demonstrates how provenance and abstention can be built into a variant-query system. These findings are promising but preliminary. Public release of the frozen data, complete run manifest, query-level annotations, and external validation is necessary before the system can support claims beyond research use.

## Data availability

The underlying records were reported to originate from public resources including CIViC, ClinVar/dbSNP, Open Targets, UniProtKB/Swiss-Prot, Ensembl, and PubMed/PMC, subject to their respective licenses and terms. The harmonized dataset, source snapshot manifest, and split assignments are available from the corresponding author upon reasonable request. To maximize reproducibility, the final preprint should deposit all redistributable processed data in a versioned public archive rather than rely only on private correspondence.

## Code availability

The reported implementation comprises automated data extraction, BioBERT fine-tuning, evidence retrieval, response generation, and a web application. Code and model artifacts are available from the corresponding author upon reasonable request.

